# Prolyl isomerase FKBP12 reduces axon growth and negatively regulates microtubule polymerization by inhibiting CRMP2A

**DOI:** 10.1101/2025.01.13.632804

**Authors:** Romana Weissova, Jan Sabo, Djamel Eddine Chafai, Jakub Ziak, Peter Buran, Satish Bodakuntla, Carsten Janke, Zdenek Lansky, Martin Balastik

## Abstract

Prolyl isomerases are enzymes catalyzing conformational change of the peptide bond between proline and the preceding amino acid, regulating the function and stability of their substrates. We have previously identified CRMP2A - the longer isoform of a microtubule-associated protein Collapsin response mediator protein 2 - as a substrate of the phospho-specific prolyl isomerase Pin1. CRMP2A is negatively regulated and destabilized by CDK5 phosphorylation in the distal axons and growth cones. Pin1 specifically binds to phosphorylated CRMP2A and stabilizes it by inducing conformational changes. However, the conformational regulation of unphosphorylated CRMP2 remains unknown. Here, we show that the prolyl isomerase FKBP12 specifically binds to unphosphorylated CRMP2A and regulates its activity. Using *in vitro* microtubule polymerization assays we demonstrate that CRMP2A promotes microtubule growth and that this function is inhibited by FKBP12. Next, using GFP-EB3 microtubule plus-end tracking assay, we demonstrate that FKBP12 inhibits CRMP2A-mediated microtubule polymerization also in cells. Furthermore, we show that FKBP12 co-localizes with unphosphorylated CRMP2A in growth cones and that expression of FKBP12 reduces axon growth in microfluidic chambers, while FKBP12 knockdown enhances it. Together, we demonstrate that FKBP12 is a negative regulator of microtubule dynamics and axon growth. Moreover, we show that two prolyl isomerases can differentially (positively or negatively) regulate activity of a common substrate depending on its phosphorylation. This provides an additional layer of phosphorylation-dependent or -independent control of protein activity, microtubule dynamics, and neuronal growth. Given the broad substrate specificity of FKBP12 and Pin1, this regulatory mechanism likely contributes to the modulation of diverse proteins and cellular processes in the nervous system and beyond.

## Introduction

Conformational changes provide a rapid mechanism to change protein properties (such as activity or stability) and represent, together with covalent posttranslational modifications, an essential level of protein regulation. While the majority of the peptide bonds are largely in *trans* conformation with regard to the position of their side chains, peptide bonds immediately preceding prolyl residues are prone to conformational changes between *trans* and *cis* conformation with a possible direct effect on the function of the protein. Conformational changes of those peptide bonds are catalyzed by peptidyl-prolyl *cis/trans* isomerases (PPIs), ubiquitous proteins expressed in both eukaryotes and prokaryotes (Schmid, 2001). Prolyl isomerases are regulators of many cellular processes such as transcription or the cell cycle (Aghdasi et al., 2001; Hanes, 2015; Lin et al., 2015; Shaw, 2007) and the deregulation of many prolyl isomerases has been linked to several human diseases such as cancer, and cardiovascular or neurodegenerative disorders (Blair et al., 2015; Gerard et al., 2011; Perrucci et al., 2015; Theuerkorn et al., 2011).

There are 3 structurally diverse families of prolyl isomerases – cyclophilins, FK506-binding proteins (FKBPs), and parvulins (Schmid, 2001). Among them, Pin1 of the parvulin family is the only known phospho-specific prolyl isomerase specifically recognizing substrates after their phosphorylation at serine or threonine immediately preceding proline (Yaffe et al., 1997). We have shown that Pin1 plays a critical role in neural development by changing the conformation and stability of one isoform of Collapsin response mediator protein 2 - CRMP2 (Balastik et al., 2015). CRMP2 is a microtubule-associated protein that is highly expressed in neurons mainly during development. CRMP2 has been shown to bind microtubules, promote their polymerization, and mediate growth cone collapse upon Semaphorin 3A signaling (Fukata et al., 2002; Uchida et al., 2005). We have demonstrated that CRMP2 is downstream of Semaphorin 3F signaling and that its deficiency results in synapse pruning defects and histological and behavioral changes associated with autism spectrum disorder (Ziak et al., 2020). As such, CRMP2 is an important regulator of neural development.

The activity of CRMP2 is regulated at multiple levels e.g. phosphorylation by kinases such as Cdk5, GSK3β, and Rho kinase has been shown to control its affinity to tubulin (Nakamura et al., 2020). Moreover, CRMP2 exists in two isoforms produced by alternative splicing. CRMP2B, a shorter isoform, is particularly abundant in the adult brain, and has been analyzed in most of the previous studies. CRMP2A is a longer isoform extended at the N-terminus by an alternative exon. Although CRMP2A is generally less expressed, it is abundant in axonal growth cones (Balastik et al., 2015; Bretin et al., 2005). The functional differences between those two isoforms are still elusive. We have shown that the CRMP2A isoform plays a unique role in distal axons, where its function and stability is regulated by Cdk5 phosphorylation and subsequent isomerization by prolyl isomerase Pin1 (Balastik et al., 2015). While Pin1-mediated isomerization demonstrates regulation of phosphorylated CRMP2A, whether (and what) prolyl isomerase controls conformation of the unphosphorylated CRMP2A and what effect on microtubules and axons it has, is not known.

Here, we demonstrate that prolyl isomerase FKBP12 (FK506 binding protein-12; FKBP1A) specifically binds to and inhibits the activity of unphosphorylated CRMP2A. FKBP12 was first described for its role in immunoregulation as a major binder of immunosuppressant FK506 (Siekierka et al., 1989). Later, it was shown that FKBP12 is highly expressed in the brain (Steiner et al., 1992) and has been linked to neurodegeneration, synaptic plasticity, or neuronal regeneration (Caminati and Procacci, 2020; Gold et al., 1997; Hoeffer et al., 2008; Lyons et al., 1995). We show that FKBP12 interacts with CRMP2A and that phosphorylation of CRMP2A abolishes the binding. Moreover, we demonstrate that FKBP12 regulates CRMP2A activity. While CRMP2A promotes microtubule polymerization *in vitro* and *in vivo*, FKBP12 significantly inhibits this function. In contrast, the de-phosphomimetic mutant of CRMP2A (that binds to FKBP12 more efficiently) does not promote microtubule polymerization, unless FKBP12 is silenced in the cell. Finally, we demonstrate that FKBP12 co-localizes with unphosphorylated CRMP2A in the growth cones and that knockdown of FKBP12 in neurons promotes axon growth in microfluidic chambers, while expression of FKBP12 reduces it.

Together, our findings reveal that conformational changes in the unphosphorylated CRMP2A catalyzed by the prolyl isomerase FKBP12 act as a negative regulator of microtubule dynamics and neural growth. Furthermore, given that the prolyl isomerase Pin1 has been shown to positively regulate axon growth both in vitro and in vivo by isomerizing phosphorylated CRMP2A, our results uncover a novel regulatory layer whereby prolyl isomerases provide positive or negative modulation of protein activity. This mechanism, alongside or in conjunction with protein phosphorylation, orchestrates microtubule dynamics, axon growth, and neural development.

## Results

### Prolyl isomerase FKBP12 binds to CRMP2A

Phosphorylated CRMP2 (isoform A) is specifically bound and regulated by prolyl isomerase Pin1 (Balastik et al., 2015). In order to understand the conformational regulation of unphosphorylated CRMP2, we first tested whether prolyl isomerase FKBP12, which shares multiple substrates with Pin1 (Blair et al., 2015), binds to CRMP2A. Using anti-CRMP2A antibodies, we were able to co-immunoprecipitate FKBP12 together with CRMP2A from WT (wild-type), but not CRMP2-KO (CRMP2 knock-out) mouse brain lysates (Figure 1A).

**Figure 1.**
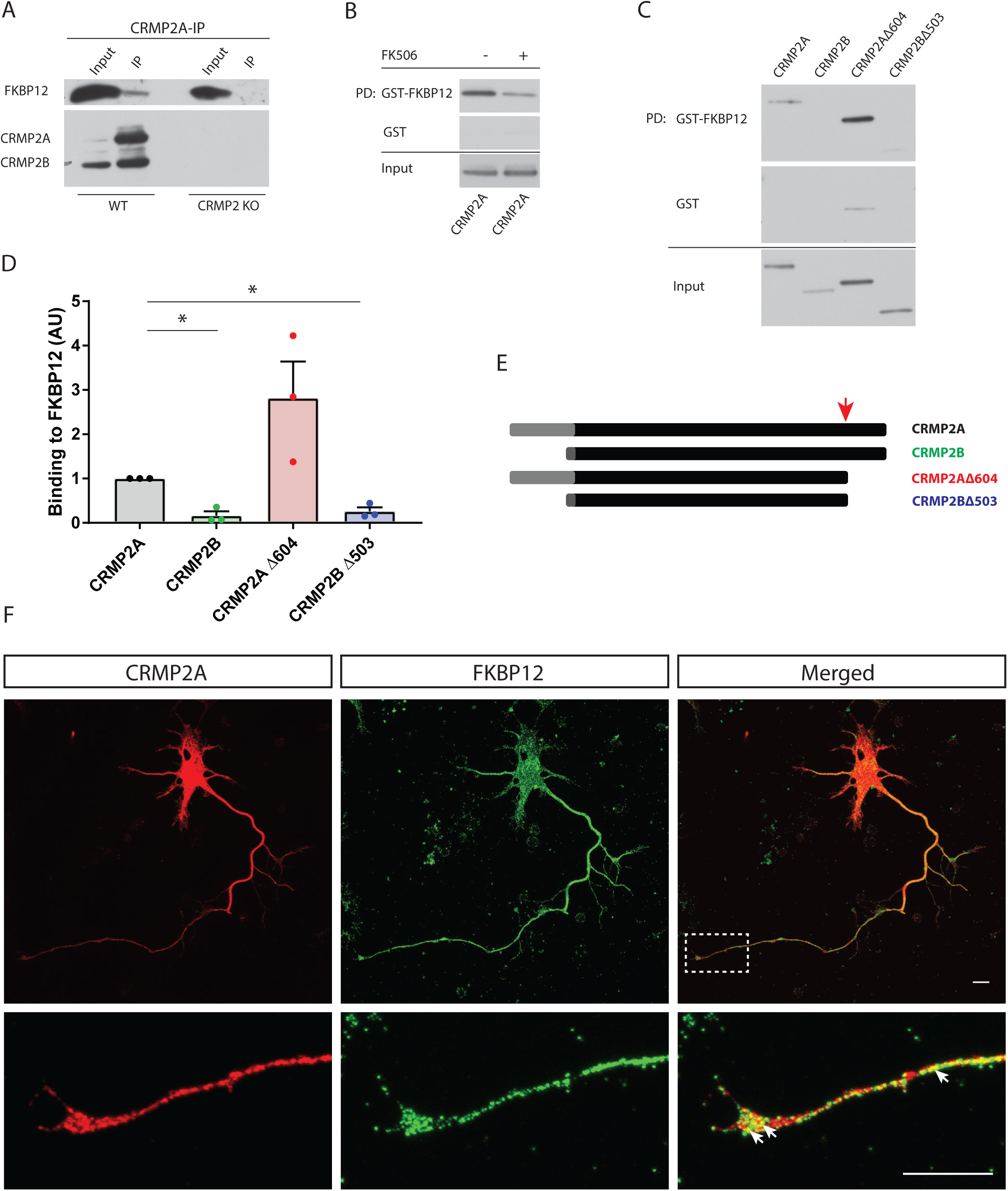
Prolyl isomerase FKBP12 interacts with CRMP2A *in vitro* and in axonal growth cones. A) FKBP12 co-immunoprecipitates with CRMP2A. Brains of young wild-type mice (WT) or CRMP2 knock-out mice (CRMP2-KO) were homogenized and used for immunoprecipitation using anti-CRMP2A antibody. FKBP12 co-immunoprecipitates with CRMP2A in WT but not in CRMP2-KO lysate which was used as a negative control. B) Pull-down of CRMP2A by GST-FKBP12 (or GST control) with or without FKBP12 inhibitor (FK506). The presence of FK506 reduces CRMP2A-FKBP12 interaction. C-E) FKBP12 binds preferably to CRMP2A isoform. C-D) FKBP12 preferentially binds to CRMP2A through its N-terminus. FKBP12 pull-down from HEK293T cells expressing CRMP2 isoforms or their truncated forms (shown in E). D) The quantification of the WB shown in C) normalized to the input (mean ± SEM, *p < 0.05). E) Schematic representation of CRMP2A and CRMP2B isoforms and their truncated forms (CRMP2AΔ604, CRMP2BΔ503) used for pull-down analysis (red arrow marking the deletion location, the alternative exon is shown in gray). F) CRMP2A and FKBP12 are both abundant in primary cortical neuron cultures, particularly in axons and their growth cones (arrows). Neurons were stained at DIV4 with antibodies against FKBP12 and CRMP2A. Scale bar 10 µm.

To further support the interaction and analyze it in more detail, we performed pull-down assays using purified GST-tagged FKBP12, and lysates of HEK293T cells expressing CRMP2 protein or its modified forms tagged with FLAG-tag (Figure 1B, C). First, to verify the specificity of the binding, we performed a pull-down in the presence or absence of FKBP12 inhibitor – FK506 (tacrolimus). The inhibitor decreased the binding distinctly, suggesting that the interaction between FKBP12 and CRMP2A is specific and competing with the inhibitor for a hydrophobic binding site pocket (Figure 1B).

Next, since CRMP2 exists in two isoforms (CRMP2A and CRMP2B; Figure 1E), and we have previously shown, that only the CRMP2A isoform is isomerized by Pin1 (Balastik et al., 2015), we tested if the binding of FKBP12 is also isoform-specific. Using pull-down assays with lysates of CRMP2A- or CRMP2B-expressing HEK293T cells, we found that, similar as Pin1, FKBP12 was preferably pulling down CRMP2A but not CRMP2B isoform (p = 0.01, Figures 1C-D) (CRMP2A = 1 ± 0; CRMP2B = 0.16 ± 0.10; CRMP2AΔ604 = 2.82 ± 0.82; CRMP2BΔ503 = 0.26 ± 0.09; mean ± SEM, Welsch’s t-test; *p < 0.05; n = 3 experiments).

CRMP2 activity is regulated by phosphorylation of several phosphorylation sites on its C-terminus, of which phosphorylation of S522 (S623 for CRMP2A isoform) is considered a priming site for further phosphorylation of the C terminus (Yoshimura et al., 2005). To test whether the C-terminus of CRMP2 regulates FKBP12 binding, we performed a pull-down assay with CRMP2A and CRMP2B lacking their C-terminal domains (Figure 1E). Deletion of the C-terminus appeared to increase (non-significantly) CRMP2A binding to FKBP12 (p = 0.16, Figures 1C-D), while it did not increase the binding of CRMP2B. Together our data indicate that FKBP12 binds to the unique N-terminus of CRMP2A (which is not present in CRMP2B) and that the C-terminal domain negatively regulates the interaction.

Both CRMP2 and FKBP12 are abundant in neurons (Kamata et al., 1998; Steiner et al., 1992). We have shown before, that CRMP2A is localized particularly in axons and growth cones (Balastik et al., 2015). To analyze whether FKBP12 and CRMP2A co-localize in neurons, we performed CRMP2A and FKBP12 double-immunostaining in primary neuron cultures. We detected a partial co-localization of FKBP12 and CRMP2A in distal axons and growth cones (Figure 1F, G) suggesting that, first, the two proteins, indeed, can interact in neurons, but, second, that their interaction is likely controlled by other mechanisms (e.g. phosphorylation) since their co-localization is only partial.

### CRMP2A phosphorylation inhibits FKBP12 binding

We have previously shown that the binding of Pin1 to CRMP2A is dependent on the phosphorylation of CRMP2A by Cdk5 (Balastik et al., 2015). To test whether FKBP12 also preferentially binds phosphorylated CRMP2A, we co-transfected HEK293T cells with CRMP2A together with Cdk5 kinase and its activator p25 and performed Pin1 or FKBP12 pull-down assays. While Cdk5 upregulation increased the binding of CRMP2A to Pin1, as shown previously ((Balastik et al., 2015); Figure 2A-B), it, surprisingly, virtually inhibited its binding to FKBP12 (Figure 2A-B), (CRMP2A = 1 ± 0; CRMP2A + Cdk5 = 0.39 ± 0.08; mean ± SEM; Welsch’s t-test; ***p = 0.0001; n = 8 experiments).

**Figure 2.**
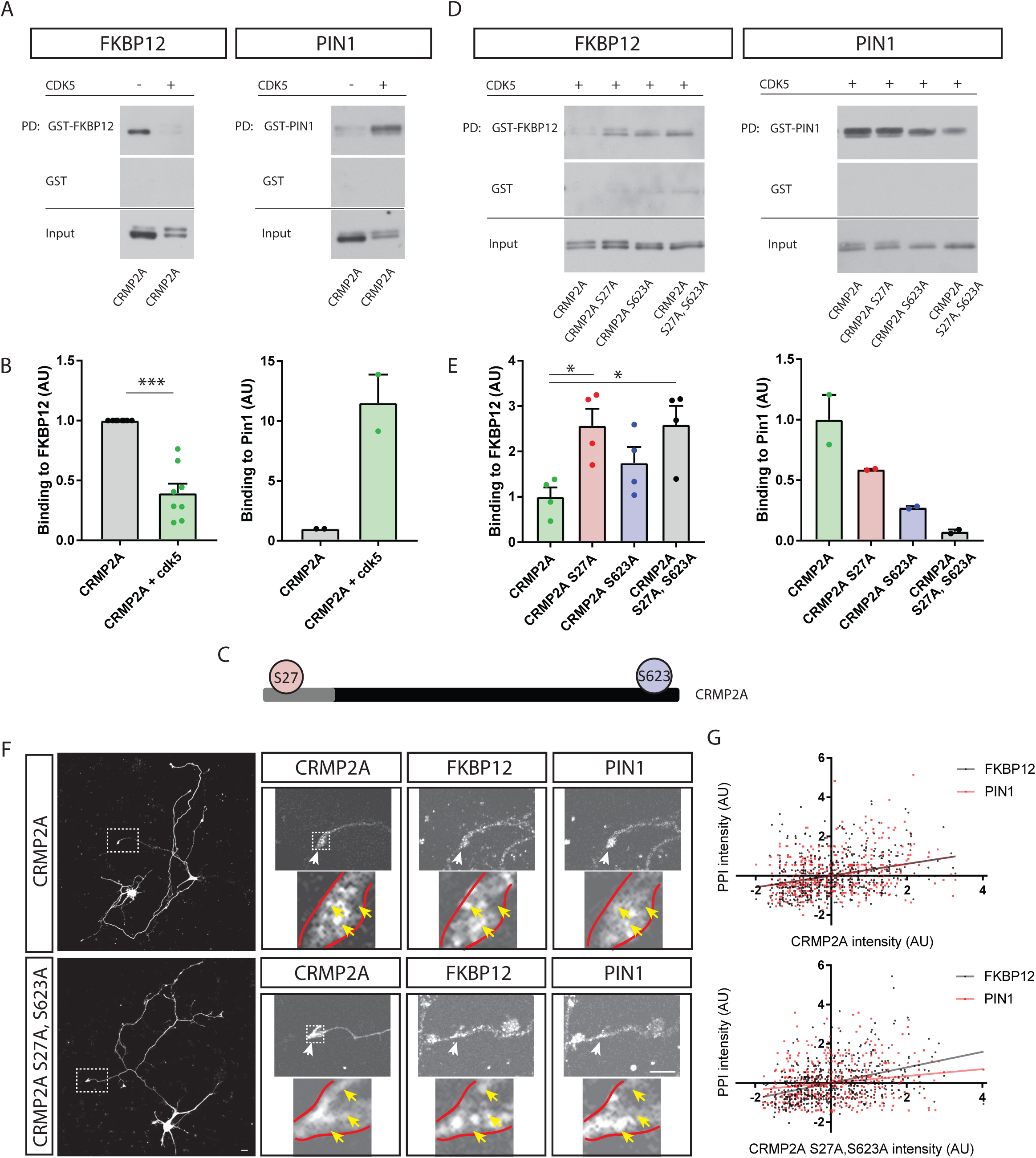
FKBP12 binds preferentially unphosphorylated, while Pin1 phosphorylated CRMP2A. A-B) FKBP12 or Pin1 pull-down of Cdk5-phosphorylated CRMP2A. Phosphorylation of CRMP2A inhibits its binding to FKBP12 but promotes its binding to Pin1. A) Representative western blots and B) quantification normalized to the input (mean ± SEM; ***p = 0.0001). C) Schematic representation of CRMP2A molecule and its two Cdk5 phosphorylation sites – S27 and S623. The alternative exon is shown in gray. D-E) Pull-down of CRMP2A and its dephospho-mimetic mutants (S27A, S623A, or S27A/S623A) by Pin1 or FKBP12 under high Cdk5 activity. S27A mutation increases the binding of FKBP12 to CRMP2A similarly to the double S27A, S623A mutant. S623A mutation shows a similar, non-significant trend. Dephospho-mimetic mutants decrease binding of CRMP2A to Pin1. E) Quantification of WB in D) (mean + SEM; *p < 0.05). F-G) Co-localization of di-dephospho-mimetic CRMP2A with FKBP12 or Pin1 in the growth cones. F) Representative images of analyzed neurons and growth cones. Scale bar 10 µm. G) Correlations between the levels of mCherry signal (corresponding to CRMP2A or its mutated variant) and levels of FKBP12 or Pin1 (immunostained), shown as scatter plots with linear regression. FKBP12 and Pin1 co-localizes (correlates) the same with CRMP2A, while FKBP12 correlates significantly more with CRMP2A S27A, S623A and Pin1 significantly less.

There are two known Cdk5 phosphorylation sites in CRMP2A, Ser27 (S27) and Ser623 (S623), out of which Ser27 is present only in the CRMP2A isoform (Figure 2C). To analyze which of the phosphorylation sites is critical for abolishing the binding to FKBP12, we performed pull-down assays using various dephospho-mimetic (Ala) mutations of the two Cdk5 sites of CRMP2A (Supplementary figure S1). Similar to what we have shown before (Balastik et al., 2015), in the presence of CDK5 the dephospho-mimetic CRMP2A mutations S27A, as well as S623A, reduced binding to Pin1 (Figure 2D-E, right panels). In contrast, the CRMP2A S27A mutation was in the presence of Cdk5 sufficient to restore its binding to FKBP12 (Figure 2D-E, left panels, p = 0.02). S623A mutation showed the same trend (but not significant) (Figure 2D-E, left panels, p = 0.1271). The combination of both mutations (S27A and S623A) increased the CRMP2A binding to FKBP12 compared to WT CRMP2A (p = 0.02), but not beyond the S27A levels (Figure 2D-E, left panels; p = 0.97) (CRMP2A = 1 ± 0.21; CRMP2A S27A = 2.57 ± 0.38; CRMP2A S623A = 1.75 ± 0.35; CRMP2A S27A, S623A = 2.59 ± 0.41; mean + SEM; Welsch’s t-test; *p < 0.05; n = 4 experiments). The data indicate that the N-terminus of CRMP2A, and its dephosphorylation, is critical for the interaction with FKBP12. The results are in accord with the data we obtained by pull-down assays using C-terminus truncated CRMP2A (Figure 1C-E) showing that removing the C-terminus of CRMP2A did not reduce FKBP12 binding.

In order to test whether CRMP2A phosphorylation controls its interaction with FKBP12 also in neurons, we analyzed the co-localization of CRMP2A (or of its di-dephospho-mimetic mutant) with FKBP12 or Pin1 within the growth cones (Figure 2F). We first transfected CRMP2A-deficient cortical neurons with vectors expressing mCherry-CRMP2A or its di-dephospho-mimetic mutant form (mCherry-CRMP2A S27A, S623A). Next, using the fluorescence, we correlated CRMP2A (or its di-dephospho-mimetic mutant) levels with endogenous (immuno-stained) FKBP12 or Pin1 (Figure 2F).

The data showed that the WT CRMP2A is co-localizing with both Pin1 and FKBP12 and that the correlation coefficients between CRMP2A - FKBP12 and CRMP2A - Pin1 in the growth cones are not significantly different (correlation coefficient r_CRMP2A-FKBP12_ = 0.2998; r_CRMP2A-Pin1_ = 0.2998; p_(between the correlations)_ = 1). Importantly, di-dephospho-mimetic mutation (mCherry-CRMP2A S27A, S623A) significantly increased co-localization of CRMP2A with FKBP12 (r = 0.3959; p_(between the correlations)_ = 0.049), but significantly decreased co-localization with Pin1 (r = 0.1763; p_(between the correlations)_ = 0.019). The co-localization of CRMP2A S27A, S623A with FKBP12 and Pin1 was thus significantly different (p_(between the correlations)_ = 0.0001). These data are in agreement with the results obtained by pull-down assays (Figure 2A-E) together demonstrating that dephosphorylation of CRMP2A (on its Cdk5 phosphorylation sites) prevents its interaction with Pin1 but promotes its binding to FKBP12 - both *in vitro* and in the neuronal growth cones.

### FKBP12 inhibits microtubule polymerization by isomerizing CRMP2A

Considering CRMP2A is a microtubule-associated protein regulating microtubule dynamics (Fukata et al., 2002), we next tested whether FKBP12 also regulates microtubule dynamics through its interaction with CRMP2A. First, we performed an *in vitro* tubulin turbidity assay. As expected, the application of CRMP2A promoted tubulin polymerization over the control tubulin sample, as assessed by measuring Abs_350_ nm in the plateau phase (Figure 3A-B; Ctrl = 0.065 ± 0.002; CRMP2A = 0.083 ± 0.005). Importantly, the addition of FKBP12 abolished the effect of CRMP2A on tubulin polymerization (Figure 3A-B, S4; CRMP2A + FKBP12 = 0.064 ± 0.002) indicating that FKBP12 is negatively regulating CRMP2A. Next, we tested whether the isomerase activity of FKBP12 is critical for the inhibition of CRMP2A. Indeed, the addition of an inhibitor of FKBP12 isomerase activity (FK506) rescued the effect of CRMP2A on tubulin polymerization indicating that CRMP2A is inhibited by conformational change catalyzed by FKBP12 (Figure 3A-B; CRMP2A + FKBP12 + FK506 = 0.080 ± 0.005). Together, the results show that CRMP2A increases the growth of microtubules *in vitro* and FKBP12 inhibits this effect.

**Figure 3.**
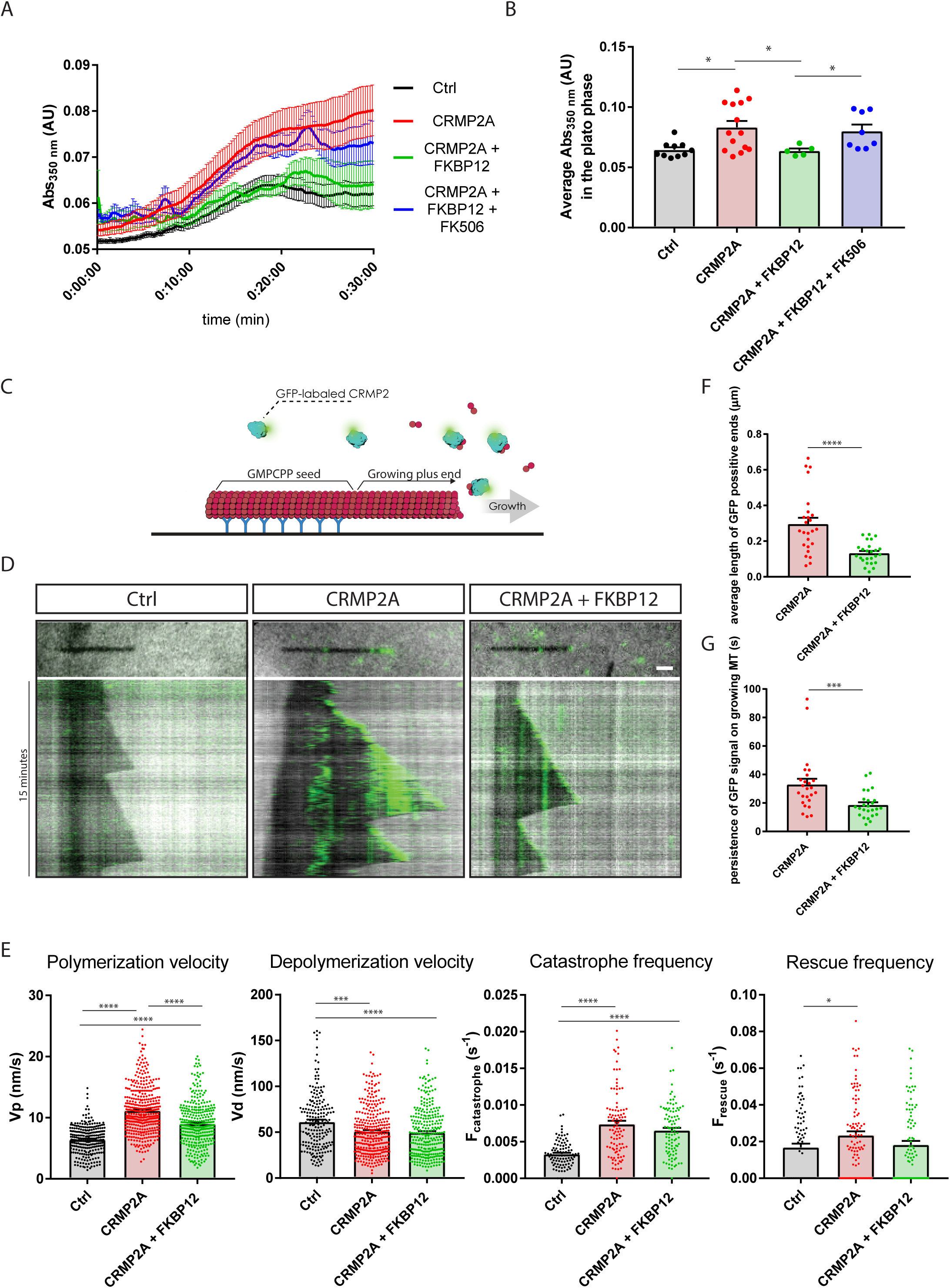
FKBP12 negatively regulates microtubule dynamics *in vitro*. A-B) CRMP2A increases microtubule polymerization in tubulin turbidimetry assay *in vitro*, and FKBP12 abolishes this effect. Application of FKBP12 together with its inhibitor (FK506) rescues the CRMP2A effect on tubulin polymerization. B) Quantification of the turbidity assay, values in the plateau phase are shown (Ctrl = 0.065 ± 0.002; CRMP2A = 0.083 ± 0.005; CRMP2A + FKBP12 = 0.064 ± 0.002; CRMP2A + FKBP12 + FK506 = 0.080 ± 0.005; mean ± SEM; Mann-Whitney test; *p < 0.05; n = 10, 14, 5, 8 experiments). C-E) CRMP2A increases microtubule polymerization in *in vitro* microtubule polymerization assay, and FKBP12 abolishes this effect. C) Schematic representation of the *in vitro* microtubule polymerization assay. D) Representative kymographs demonstrating the effect of CRMP2A and FKBP12 on microtubule polymerization (dark – microtubules; green – GFP-CRMP2A). E) Quantification of the microtubule dynamics parameters. CRMP2A increases the microtubule polymerization rate, decreases the depolymerization rate, and increases the frequency of both catastrophes and rescues. FKBP12 inhibits the effect of CRMP2A mainly by reducing the polymerization rate (shown means ± SEM; Mann-Whitney tests: *p < 0.05, ***p < 0.001, ****p < 0.0001; Polymerization velocity: n = 259, 401, 350 polymerization events in 5 independent experiments; Depolymerization velocity: n = 200, 326, 294 depolymerization events in 5 independent experiments; Catastrophe frequency: n = 97, 100, 98 microtubules in 5 independent experiments; Rescue frequency: n = 95, 99, 94 microtubules in 5 independent experiments). F) Quantification of average length of GFP-positive ends (shown means ± SEM; ****p<0.0001; Mann-Whitney test; n = 25 microtubules in 5 independent experiments). G) Quantification of the persistence of GFP signal on the growing microtubules (shown means ± SEM; ***p<0.001; Mann-Whitney test; n = 25 microtubules in 5 independent experiments). Scale bar 1 µm.

The application of FKBP12 resulted in differences in both the slope of the curves in the exponential phase of tubulin polymerization as well as their plateau phase (Figure 3A). This suggests that FKBP12 could affect tubulin polymerization rate, microtubule rescue frequencies, or both. To distinguish the two possible mechanisms, we performed *in vitro* tubulin polymerization assays using TIRF microscopy (Figure 3C). In accordance with the results of the turbidity assay, GFP-CRMP2A bound to growing microtubules and increased microtubule polymerization rate (ctrl = 6.45 ± 0.14 nm/s; CRMP2A = 11.17 ± 0.19 nm/s; p_Mann-Whitney_ < 0.0001), decreased microtubule depolymerization rate (ctrl = 61.5 ± 2.2; CRMP2A = 51.0 ± 1.4; p_Mann-Whitney_ = 0.0001) and increased the frequency of both catastrophes (ctrl = 0.0033 ± 0.0002 s^-1^; CRMP2A = 0.0074 ± 0.0004 s^-1^; p_Mann-Whitney_ < 0.0001) and rescues (ctrl = 0.017 ± 0.002 s^-1^; CRMP2A = 0.023 ± 0.002 s^-1^, p_Mann-Whitney_ = 0.0199) (Figure 3D-E). Adding FKBP12 significantly reduced the CRMP2A-mediated microtubule polymerization rate (Polymerization velocity: CRMP2A + FKBP12 = 9.02 ± 0.19 nm/s; p_Mann-Whitney_ < 0.0001), while it had no significant effect on microtubule depolymerization velocity (CRMP2A + FKBP12 = 49.7 ± 1.6 nm/s, p_Mann-Whitney_ > 0.05) (Figure 3D-E). Microtubule rescue frequency also showed a decreasing trend upon addition of FKBP12, although not significant (Rescue rate: CRMP2A + FKBP12 = 0.018 ± 0.002 s^-1^; p = 0.058; Catastrophe rate: CRMP2A + FKBP12 = 0.0066 ± 0.0003 s^-1^; p_Mann-Whitney_ > 0.05) (Figure 3D-E). Thus, FKBP12 negatively regulates microtubule dynamics by reducing particularly the tubulin polymerization rate. Interestingly, while GFP-CRMP2A forms large comets at the microtubule plus ends (Figure 3D), in the presence of FKBP12, CRMP2A appears to bind only to the very tip of growing microtubules (Figure 3D). This may indicate that FKBP12 reduces the affinity of CRMP2A to tubulin. To explore this hypothesis, we measured the average length of GFP-positive growing ends and the persistence of GFP signal on the microtubule. Both parameters decreased in the presence of FKBP12 (length of the GFP-positive ends: CRMP2A = 0.2966 ± 0.03428 um, CRMP2A+FKBP12 = 0.1329 ± 0.01207 um; mean ± SEM; pMann-Whitney <0.0001; persistence of GFP signal: CRMP2A = 33.07 ± 3.955 s, CRMP2A+FKBP12 = 18.69 ± 1.844 s; pMann-Whitney =0.0005) (Figure 3F-G). Together, these results show that FKBP12 is inhibiting CRMP2A function by lowering the binding of CRMP2A to microtubules.

### FKBP12 negatively regulates microtubule growth in cells

To test whether FKBP12 negatively regulates microtubule dynamics not only *in vitro* but also in cells, we visualized microtubule dynamics in IMCD3 cells using GFP-tagged microtubule plus end-tracking protein EB3 (Figure 4A). GFP-EB3 was expressed in the IMCD3 cells together with CRMP2A and FKBP12, or FKBP12 was silenced by shRNA (Supplementary figure S5A, B). We then analyzed the average velocity of growing microtubules. In agreement with the *in vitro* data (Figure 3), the expression of CRMP2A significantly increased the velocity of microtubule growing ends (p < 0.0001; Figure 4B-C). Co-expression of FKBP12 reverted the CRMP2A-increased velocity significantly (p < 0.0001) almost to the control level (p = 0.016; Figure 4B-C), while FKBP12 knockdown had no significant effect (p = 0.430). These results are again in agreement with the results of the *in vitro* microtubule polymerization assays (Figure 3). Importantly, unlike CRMP2A, expression of dephospho-mimetic mutant CRMP2A-S27A (which binds stronger to FKBP12 even upon Cdk5 activation) (Figure 2D), did not increase microtubule growth velocity (p = 0.2), possibly due to increased binding and inhibition by FKBP12 (Figure 4B-C). Indeed, simultaneous knockdown of FKBP12 significantly increased microtubule growth rate upon CRMP2A-S27A expression (p < 0.0001) – i.e. to a similar level as CRMP2A (p = 0.822). Co-expression of CRMP2A-S27A with FKBP12 had no significant effect on microtubule growth (p = 0.994) (Figure 4B-C) indicating that the endogenous level of FKBP12 was high enough to inhibit CRMP2A-S27A. (relative MT growth values: ctrl = 1 ± 0.01; FKBP12 = 1.01 ± 0.02; shFKBP12 = 1.03 ± 0.02; CRMP2A = 1.13 ± 0.01; CRMP2A + FKBP12 = 1.06 ± 0.02; CRMP2A shFKBP12 = 1.11 ± 0.02; CRMP2A S27A = 1.01 ± 0.01; CRMP2A S27A + FKBP12 = 1.02 ± 0.02; CRMP2A S27A shFKBP12 = 1.12 ± 0.02). Together, the data demonstrate that FKBP12 expression inhibits the CRMP2A-mediated microtubule polymerization also in cells.

**Figure 4.**
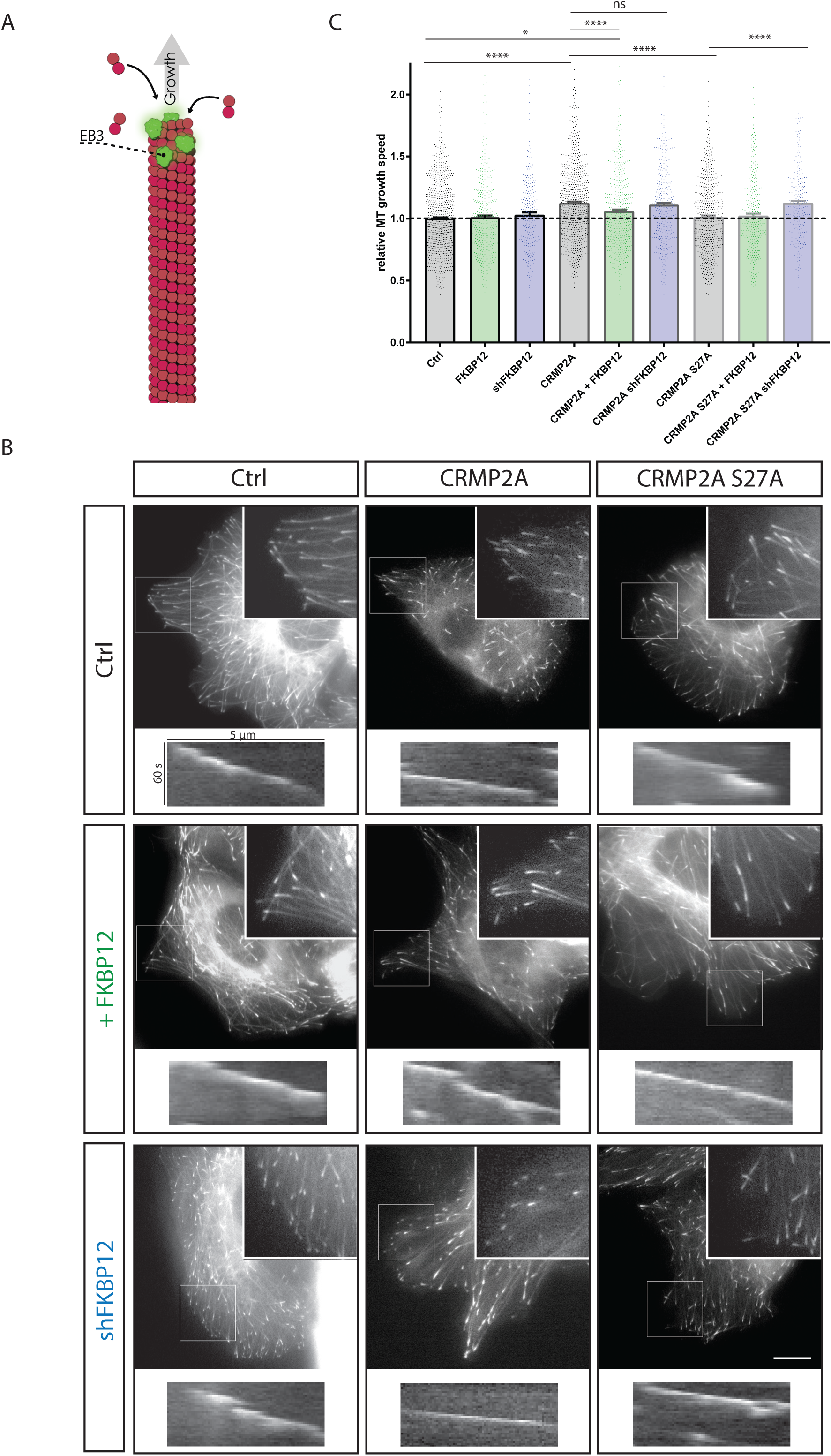
FKBP12 negatively regulates microtubule growth in cells. GFP-EB3 was used to track the dynamics of growing microtubule ends in cells. A) Schematic representation of GFP-EB3 binding to growing ends of microtubules. B-C) FKBP12 inhibits CRMP2A-mediated microtubule growth. GFP-EB3 and CRMP2A (or CRMP2A S27A) were expressed in IMCD3 cells together with an FKBP12 expressing or silencing vector. CRMP2A expression significantly increases microtubule growth velocity, but its effect can be abolished by the expression of FKBP12. CRMP2A S27A (stronger binding to FKBP12) increases microtubule growth only when combined with knockdown of FKBP12. Representative images and kymographs of analyzed microtubules shown in B). C) Quantification of relative microtubule growth rate (shown mean ± SEM; Mann-Whitney test; ns = not significant, **p < 0.01, ***p < 0.001, ****p < 0.0001; n = 761, 390, 235, 746, 462, 330, 550, 300, 235 microtubules in at least 7 independent experiments). Scale bars 10 µm.

### The level of FKBP12 in neurons modulates axon growth

CRMP2 and its isoform CRMP2A have been shown to promote axon growth and to regulate axon guidance in the developing neurons (Balastik et al., 2015; Fukata et al., 2002; Ziak et al., 2020). Similarly, inhibition of microtubule polymerization has been demonstrated to negatively affect axon growth (Tanaka et al., 1995). Our data demonstrate that FKBP12 reduces microtubule polymerization by inhibiting CRMP2A activity. Moreover, we show that FKBP12 co-localizatizes with un-phosphorylated CRMP2A in the axonal growth cones (Figure 2F, G) indicating that FKBP12 levels in the developing neurons could regulate axonal growth. To test this hypothesis, we cultured primary cortical neurons in microfluidic chambers (Figure 5A) and quantified the average axonal growth rate upon modulating FKBP12 levels. Knockdown of FKBP12 (shFKBP12) significantly increased axonal growth (ctrl = 6.41 ± 0.15 nm/s, shFKBP12 = 6.94 ± 0.18 nm/s; p_Mann-Whitney_ = 0.04; Figure 5B-D). In contrast, expression of FKBP12 significantly decreased the axon growth rate (+FKBP12 = 5.83 ± 0.16 nm/s, p_Mann-Whitney_ = 0.007; Figure 5B-D). These results are in agreement with the effect of FKBP12 on microtubules, which we measured in the IMCD3 cells, as well as *in vitro* using microtubule polymerization assays, and together identify FKBP12 as a new negative regulator of microtubule polymerization and axon growth.

**Figure 5.**
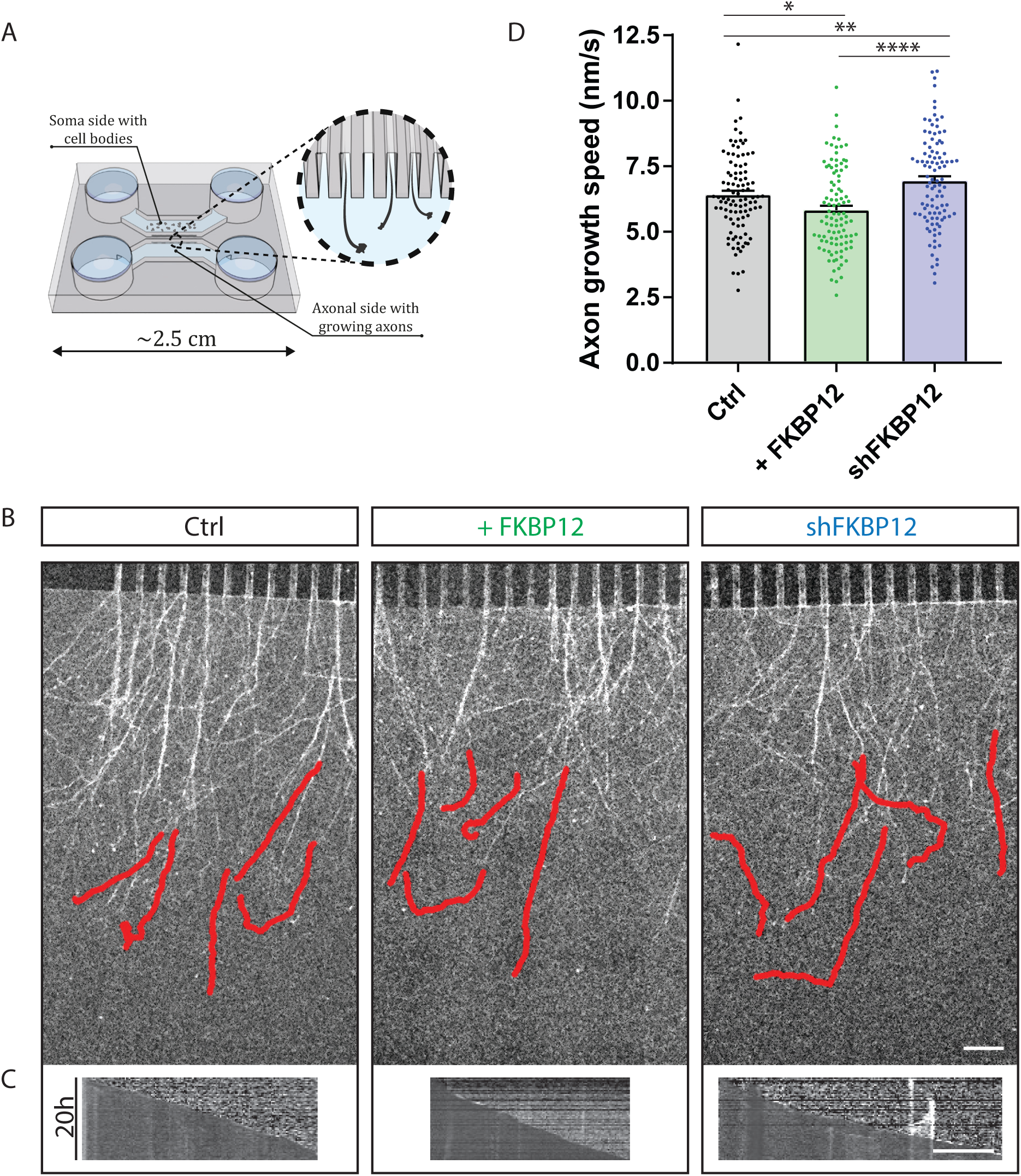
FKBP12 modulates axonal growth in cultured neurons. A) Schematic representation of microfluidic chambers used in the experiments. B) Primary embryonic cortical neurons were grown in microfluidic chambers, visualized by Cholera toxin Subunit B, and their growth rate was analyzed. Knockdown of FKBP12 (shFKBP12) increases axon growth rate while expression of FKBP12 (+FKBP12) decreases it. B) Representative images with the trajectory of axons growing for 20 hours. C) representative kymographs corresponding to the trajectories. D) Quantification of average axon growth rate in nm/s (shown means ± SEM; Mann-Whitney test; *p < 0.05, **p < 0.01, ****p < 0.0001; n=100 axons in 4 independent experiments). Scale bars 100 µm, 10h.

## Discussion

During brain development, microtubule-associated protein CRMP2 either promotes microtubule polymerization and axon growth or mediates Semaphorin 3A-induced growth cone collapse and axon retraction (Uchida et al., 2005). The dual function of CRMP2 is tightly controlled by several mechanisms. Previously, phosphorylation has been shown to regulate CRMP2 through a competing activity of protein kinases and phosphatases. While upregulation of various kinases (e.g. Cdk5 or GSK3β) has been shown to increase phosphorylation of CRMP2, inhibit its activity and axon growth (Crews et al., 2011; Uchida et al., 2005; Yoshimura et al., 2005) upregulation of phosphatases (e.g. PP2A) has been shown to promote axon growth by dephosphorylating CRMP2 (Zhu et al., 2010). In addition, conformational changes are considered to control CRMP2 activity, but its impact and mechanism of action, has so far been largely unknown. We have previously demonstrated that conformational changes of (specifically) Cdk5-phosphorylated CRMP2A isoform catalyzed by prolyl isomerase Pin1 counteracts the inhibitory effect of CRMP2A phosphorylation (Balastik et al., 2015). This allows Pin1-containing axons to withstand low concentrations of repulsive guidance cue Semaphorin 3A in the environment (which activates Cdk5) and enables these axons to grow further into the Semaphorin3A gradients than Pin1-deficient axons. Whether the unphosphorylated CRMP2 is conformationally regulated has not been shown by now. Here, we provided evidence that unphosphorylated CRMP2 – again specifically the CRMP2A isoform - is conformationally regulated by another prolyl isomerase - FKBP12. In contrast to Pin1, which stabilizes CRMP2A and promotes axonal growth (Balastik et al., 2015), FKBP12 inhibits the activity of CRMP2A, which results in lower microtubule polymerization activity *in vitro* and *in vivo*, binds to CRMP2A in growth cones and reduces axon growth. Our data thus demonstrate that prolyl isomerases are not merely maintaining native/active conformation of their substrates, but that they can form a well-orchestrated, positive or negative regulatory system, that, in line with protein phosphorylation, modulates activity of their substrates with direct biological consequences.

### CRMP2A as a specific target for conformational regulation by prolyl isomerases

CRMP2 exists in two isoforms produced by alternative splicing. The shorter, more abundant CRMP2B isoform has been so far more analyzed and shown to promote microtubule growth (Fukata et al., 2002). We now demonstrate that CRMP2A also promotes microtubule growth *in vitro* and *in vivo*. CRMP2A has an N-terminal domain which is not present in CRMP2B and it contains a unique Cdk5-phosphorylation site (S27) (Balastik et al., 2015). Phosphorylation of this site and another C-terminal Cdk5 phosphorylation site (S623 on CRMP2A, corresponding to S522 on CRMP2B) is required for binding to prolyl isomerase Pin1 (Balastik et al., 2015). Here we show that particularly S27 is important for binding to prolyl isomerase FKBP12 and that its phosphorylation serves as a switch from isomerization by FKBP12 to Pin1. The critical role of the N-terminal S27 for the conformational regulation of CRMP2 also explains why only the longer CRMP2A isoform is regulated by prolyl isomerases.

Previously, phosphorylation of CRMP2 has been shown to inhibit its function (Uchida et al., 2005). Interestingly, our data indicate that even dephosphorylated CRMP2 - specifically isoform A - may be readily inactivated by FKBP12 isomerization. Indeed, dephospho-mimetic CRMP2A mutant (CRMP2A-S27A) was not able to promote the microtubule growth in cells (Figure 4B, C) - likely due to its higher affinity to FKBP12 (Figure 2D, E) and subsequent conformational inhibition. Supporting this, upon FKBP12 knockdown, CRMP2A-S27A was able to promote microtubule growth (Figure 4B, C), and overexpression of FKBP12 inhibited microtubule growth promotion induced even by WT CRMP2A (Figure 4B, C).

CRMP2A isoform is less abundant in the adult brain, but its relative amount is higher in active growth cones during development (Balastik et al., 2015), when precise regulation of microtubule growth and guidance is critical. Thus, a combination of various levels of phosphorylation, alternative splicing, and isomerization of CRMP2A could help to diversify responses of different growing axons to extracellular stimuli, which is necessary for the formation of specific neuronal connectivity pattern.

### FKBP12 as a microtubule regulator in development and disease

FKBP protein family contains in humans at least 15 members (Tong and Jiang, 2016) targeting several cytoskeleton-related substrates such as microtubule-associated protein tau isomerized by FKBP12, FKBP51, FKBP52 (Chambraud et al., 2010; Ikura and Ito, 2013; Jiang et al., 2023; Jinwal et al., 2010). Importantly, two FKBP family members – FKBP52 and FKBP25 – have been previously shown to regulate microtubule polymerization directly by binding to tubulin (Chambraud et al., 2007; Dilworth et al., 2018). FKBP12 was considered not to regulate microtubule polymerization since (as one of the smallest FKBP family members) it has no tubulin binding domain and no direct effect on tubulin polymerization (Chambraud et al., 2007). In contrast, we demonstrate here, that FKBP12 also controls microtubule polymerization, but its effect is indirect by isomerization of CRMP2A protein.

FKBP12 has been studied mainly in connection to immunoregulation and cardiac research, but its role in neurons and brain function has been also analyzed. It was shown that expression of FKBP12 is enhanced during neural regeneration (Lyons et al., 1995; Mason et al., 2003) and at the same time, it was shown, that FKBP12 inhibitor (FK506) as well as another inhibitor without immunosupressing function accelerates nerve regeneration (Gold et al., 1995; Gold et al., 1997; Khan et al., 2002). Moreover, FK506 has been shown to promote neurite outgrowth (Lyons et al., 1994), which is all in agreement with our in vitro and in vivo data. The mechanism by which FK506 inhibitor supports neurite growth and nerve regeneration has not been well understood, but our data indicate that activation of CRMP2A could play an important role. Indeed, CRMP2 was shown to promote growth of axons, its mRNA level increases upon hypoglossal nerve injury and it promotes its regeneration (Suzuki et al., 2003). Similarly, moderate stabilization of microtubules with microtubule-stabilizing drugs has been demonstrated to enhance axon regeneration and growth (Hellal et al., 2011; Ruschel et al., 2015; Sengottuvel et al., 2011).

In mouse models, brain-specific knockout of FKBP12 has been shown to result in enhanced late-phase long-term potentiation (L-LTP) and perseveration in several memory assays (Hoeffer et al., 2008). In contrast, brain-specific CRMP2 deletion leads to reduced LTP and impaired learning and memory (Zhang et al., 2016; Ziak et al., 2020), which is again in accord with the inhibitory role of FKBP12 on CRMP2. Deregulation of both FKBP12 as well as CRMP2 was also linked to behavioral changes associated with neurodevelopmental disorders, including autism spectrum disorder (ASD), obsessive-compulsive disorder (OCD), and schizophrenia (Hoeffer et al., 2008; Zhang et al., 2016; Ziak et al., 2020).

Prolyl isomerases are becoming the center of attention not only because of their roles in physiological conditions as signaling regulators, but they were also connected to many diseases mainly so-called conformational diseases such as Alzheimer’s disease. Moreover, it was shown that both CRMP2 and FKBP12 accumulate in neurofibrillary tangles (Sugata et al., 2009; Yoshida et al., 1998). Another microtubule-associated protein and marker of neurofibrillary tangles - tau - is bound by prolyl isomerases FKBP12 and Pin1 and it was shown that both are regulators of tau aggregation into neurofibrillary tangles (Ikura and Ito, 2013; Kimura et al., 2013).

The data obtained in this and previous studies (Balastik et al., 2015) together propose a new mechanism of conformational, isoform-specific regulation of CRMP2 by two prolyl isomerases, Pin1 and FKBP12, controlling neuron growth (Figure 6). In distal axons, the growth of microtubules is supported by CRMP2A that binds to tubulin heterodimers and promotes microtubule polymerization (Figure 6A). The levels of CRMP2A are negatively regulated by CDK5, that phosphorylates and destabilizes CRMP2A, leading to its degradation in proteasome and reduction of axon growth (Balastik et al., 2015) (Figure 6B). At the same time, the CDK5-phosphorylated CRMP2A becomes a substrate of Pin1 that catalyzes its conformational change that stabilizes the protein allowing it to be dephosphorylated by protein phosphatases and, in the native conformation, to promote microtubule and axonal growth. The activity of dephosphorylated CRMP2A is then negatively regulated by FKBP12, that specifically binds to it, inhibits its microtubule-polymerization activity and reduces axon growth (Figure 6C).

**Figure 6.**
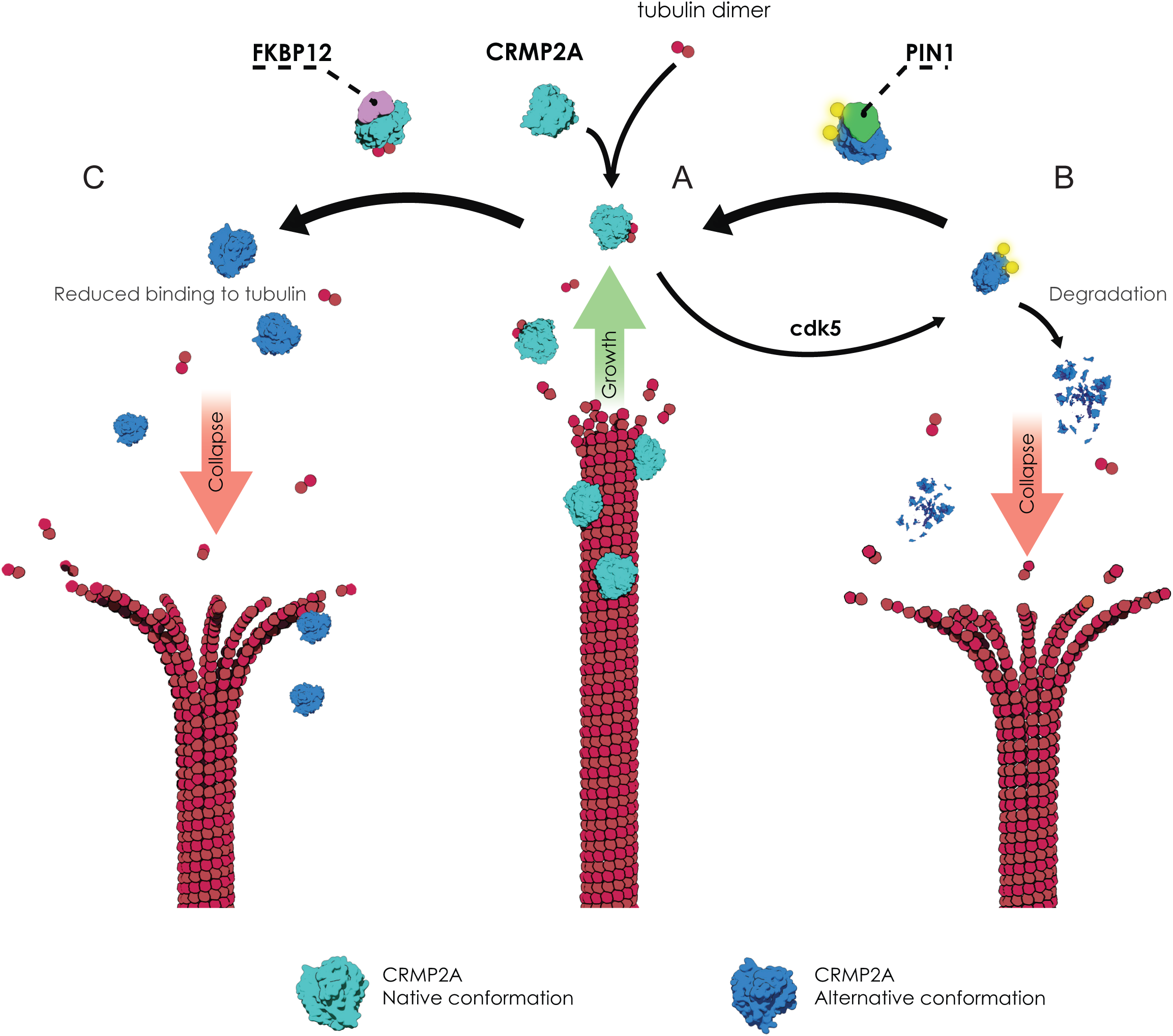
Model of CRMP2A conformational regulation and its effect on microtubule dynamics and axon growth. A) CRMP2A binds to tubulin heterodimers and promotes microtubule polymerization and axon growth. B) Phosphorylation by CDK5 destabilizes CRMP2A in distal axons leading to its degradation in proteasome and reduction of axon growth. CDK5-phosphorylated CRMP2A is bound by Pin1 that stabilizes the protein and promotes microtubule polymerization and axonal growth. C) The dephosphorylated CRMP2A is in growth cones bound and negatively regulated by FKBP12, which inhibits microtubule-polymerization and reduces axon growth.

Interestingly, both Pin1 and FKBP12 target the same two Cdk5-dependent phosphorylation sites of CRMP2A. In the case of Pin1, the two phosphorylation sites seem to contribute equally, while for FKBP12 the isomerization of the N-terminal site (Ser27) of CRMP2A seems to be critical. This may be due to Pin1 containing two domains capable of binding phospho-Ser/Thr-Pro motifs in its substrates (WW and PPIase domains), while only one (PPIase) domain is present in FKBP12.

In conclusion, our study uncovers a novel molecular system in which two distinct prolyl isomerases regulate the function of a common substrate in opposing ways, depending on its phosphorylation state. Furthermore, our findings highlight how the interplay between different prolyl isomerases and specific kinases can fine-tune microtubule dynamics in growth cones, a process critical for precise axon growth and proper brain development.

## Methods

### Plasmids

Plasmids for expression of mouse CRMP2A and CRMP2B with N-terminal FLAG-tag in pCDNA3, as well as mutated forms of CRMP2A (S27A, S623A, and S27A+S623A), were published before (Balastik et al., 2015). Sequences of truncated forms of CRMP2A (1-604) and CRMP2B (1-503) were prepared using PCR from full-length CRMP2A or CRMP2B and cloned into pCDNA3 with N-terminal FLAG-tag. For the purification of GST-tagged proteins from bacteria, pGEX-KG-GST and pGEX-KG-GST-Pin1 were used (Balastik et al., 2015). FKBP12 sequence was prepared by PCR from mouse brain cDNA library and inserted into pGEX-KG-GST to produce fused protein GST-FKBP12.

FLAG-tagged CRMP2A plasmid together with pCDNA3-Cdk5 and pCDNA3-p25 (Balastik et al., 2015; Patrick et al., 1999) were used for phosphorylation of CRMP2A by CDK5 in cells. Fusion mCherry-CRMP2A was prepared by insertion of mCherry in frame, 5’ of CRMP2A sequence in pCDNA3.1 vector. Mutated mCherry-CRMP2A (S27A) was prepared by site-directed mutagenesis as described previously (Balastik et al., 2015).

For purification of CRMP2A for *in vitro* experiments, we inserted FLAG-tag 5’ of GFP-CRMP2A fusion protein in pCDNA3.1 vector.

Microtubule +tip tracking was done using a lentiviral pTRIP vector containing GFP-EB3 and CRMP2A under the CAG promoter and separated by P2A site and shRNA (either control shRNA for luciferase or shRNA for FKBP12) under the H1 promoter. For FKBP12 overexpression, the CRMP2A sequence in pTRIP was exchanged by the FKBP12 sequence using the SLIC cloning method (Jeong et al., 2012). For overexpression of FKBP12 in microfluidic chambers, we used a pTRIP vector derived from the one with GFP-EB3 and FKBP12 by removing the EB3 sequence.

FKBP12 knockdown was made by lentiviral transduction using vector pGhU6 (Radomska et al., 2012) or pGhU6 with marker GFP fluorescent protein exchanged for mCherry. The target sequence for shRNA design was selected using the TRC library database (TRCN0000012492 – TGATTCCTCTCGGGACAGAAA; nonspecific control (NSC) - ATCTCGCTTGGGCGAGAGTAAG).

### All insertions were confirmed by DNA sequencing

#### Animals

All experimental procedures were performed in compliance with European directive 2010/63/EU and were approved by the Czech Central Commission for Animal Welfare. Mice (C57BL6/N background) were housed and handled according to the institutional committee guidelines with free access to food and water. For immunoprecipitation, WT or CRMP2-KO murine brains were used (Ziak et al., 2020). For primary neurons culture, WT or CRMP2A-KO mice were used (Ziak et al., in preparation).

#### Antibodies

We used Anti-FLAG (Millipore, F7425; 1:5000), Anti-FKBP12 (Abcam, ab2918; 1:10000), Anti-CRMP2 (FUJIFILM Wako chemicals, 9F; 1:5000), Anti-CRMP2A ((Balastik et al., 2015); 1:10000), Anti-CRMP2A pS27 (antibody produced in rabbit using the peptide CNLGSG(Sp)PKPRQK and affinity purified using the same peptide, 1:20000), and Anti-GAPDH (Merck, G9545; 1:20000) for western blot analysis. For immunocytochemistry, we used Anti-CRMP2A ((Balastik et al., 2015), affinity purified; 1:50), Anti-FKBP12 (Abcam, ab2918, ab58072; 1:600), Anti-Pin1 (R&D systems, MAB2294; 1:250), and fluorescently labeled secondary antibodies anti-Rabbit A488, anti-Mouse A647, anti-Rabbit A594, and anti-Mouse A488 (A-11029, A21236, A-11037, A-11029, ThermoFisher Scientific; 1:500). Immunoprecipitation was done using affinity purified Anti-CRMP2A (Balastik et al., 2015).

#### Immunoprecipitation

Immunoprecipitation was performed as described (Balastik et al., 2015). Briefly, one half of the murine brain (WT or CRMP2-KO, 1-3 months old) was homogenized in lysis buffer (50 mM HEPES pH 7.4, 150 mM NaCl, 10% glycerol, 1% Triton X-100, 1.5 mM MgCl_2_, 1 mM EGTA, 100 mM NaF, 1 mM Na_3_VO_4_, 1 mM DTT, cOmplete™ EDTA-free Protease Inhibitor Cocktail (Merck)) using ultraturrax. The lysate was sonicated (30 s, 30%), centrifuged (20 minutes, 20000 x g, 4°C), and incubated with Protein A Sepharose CL-4B (Merck) for 1 hour, 4°C (pre-clearing). The sepharose beads were spun down and the lysate was incubated with 25 µl of Protein A Sepharose CL-4B (preincubated ON with antibodies (Anti-CRMP2A antibody affinity purified (Balastik et al., 2015)). After washing (4x), the beads were boiled in 2x Sample Buffer and the supernatant separate on 15%, or 7.5% SDS PAGE for western blot analysis. Proteins were visualized using antibodies anti-FKBP12 and anti-CRMP2.

#### Bacterial protein production and purification

Protein production using the pGEX-KG vector and E. Coli BL21 was done as described previously (Harper and Speicher, 2011). Glutathione-agarose beads (Millipore, G4510) were used for protein purification as described previously (Balastik et al., 2015). Briefly, bacterial pellet was resuspended in lysis buffer (50 mM TrisCl pH 8.0, 500 mM NaCl, 1 mM EDTA, 1 mM EGTA, 5% glycerol, 1 mM DTT, 1% Triton X-100, cOmplete™ EDTA-free Protease Inhibitor Cocktail (Merck)) and incubated with lysozyme (final concentration 0.2 mg/ml; Merck, L6876). The lysate was sonicated (5 x 1 minute) and centrifuged (28000 x g, 60 minutes). The supernatant was incubated with glutathione-agarose beads for 3 hours, washed, and eluted using 30 mM glutathione (Merck, G4251). Eluted protein was dialyzed (dialysis buffer: 25 mM Tris pH 7.4, 150 mM NaCl, 5% glycerol, 1 mM DTT, 50 µg/ml PMSF; SnakeSkin dialysis tubing, 10K MWCO, ThermoFisher Scientific), aliquoted and flash-frozen. Purified protein was analyzed by acrylamide gel electrophoresis (Supplementary figure S3), and the concentration of the protein was determined using Bradford assay and/or measured by NanoDrop One (ThermoFisher Scientific).

#### Peptidyl-prolyl *cis-trans* isomerase assay

PPI-ase activity was assayed in a chymotrypsin-coupled assay as described previously (Fischer et al., 1984). The activity was measured as the change in absorption at 390 nm (every 3 s for 5 minutes) due to digestion of the *trans* isoform of the peptide Suc-AAPF-pNA (Santa Cruz; 3 mM, dissolved in 20 mg/ml LiCl in trifluoroethanol) by chymotrypsin (Merck; 60 mg/ml in 1 mM HCl) in 96well plates with glass bottom (Invitrogen, M33089) in final volume 200 µl (buffer: 50 mM HEPES, 100 mM NaCl, pH8) using BioTek Cytation3. (Supplementary Figure S4)

#### Transfection and lentiviral particles production

HEK293T cells were transfected using linear polyethylenimine (PEI; Polysciences, Inc.; 23966) and cells were collected 48 hours after transfection unless stated otherwise.

Lentiviral particles were produced by co-transfection of HEK293T cells with lentiviral vectors carrying the sequence coding protein of interest or shRNA (pTRIP or pGhU6 respectively) together with Gag/Pol and VSV-G vector (1: 0.9: 0.1, (Balastik et al., 2015)). Medium containing lentiviral particles was collected 48 hours after transfection, filtered (0.45 µm pores), and used for transduction or stored at -80°C.

#### Pull-down assays

Pull-down was performed as described previously (Balastik et al., 2015). Briefly, HEK293T cells were transfected with a pCDNA3 vector expressing proteins of interest with FLAG-tag. Two days after the transfection, cells were harvested and homogenized in lysis buffer (50 mM HEPES pH 7.4, 150 mM NaCl, 10% glycerol, 0.5% Triton X-100, 1.5 mM MgCl_2_, 1 mM EGTA, 1 mM NaF, 1 mM Na_3_VO_4_, 1 mM DTT, cOmplete™ EDTA-free Protease Inhibitor Cocktail (Merck)).

Dialyzed protein fused to GST (1 µM) was incubated with the glutathione-agarose for 3 hours, washed 2x, and then incubated with 200 µl of the lysate for 4 h, 4°C, rotating. Beads were washed 5x, boiled in the 2x Sample buffer and the supernatant was subjected to western blot analysis. The bands were visualized using anti-FLAG antibodies.

In the case of inhibitor treatment, FK506 was preincubated with the protein bound on the beads for 1 hour (20 µM; InvivoGen, tlrl-fk5) before incubation with the lysate.

#### Purification of proteins produced in HEK293T cells

FLAG-tagged proteins were purified from transfected HEK293T cells. Harvested cells were resuspended in lysis buffer and sonicated (Lysis buffer: BRB40 + 0.1% Tween, 5% glycerol, 1 mM DTT, 10 µg/ml Cytochalasin D, Benzonase (5U; Merck, 70664), cOmplete™ EDTA-free Protease Inhibitor Cocktail (Merck) + 10 mM ATP). After centrifugation, the supernatant was incubated with anti-FLAG agarose beads ON, 4°C, rotating (Anti-DYKDDDDK Tag (L5) Affinity Gel Antibody; BioLegend). After washing, the protein was eluted using 3x DYKDDDDK Peptide (ThermoFisher Scientific, A36805), dialyzed against BRB80, 1 mM DTT, and flash-frozen. Purified protein was subjected to SDS-PAGE electrophoresis and stained using coomassie brilliant blue staining (Supplementary figure S2).

#### *In vitro* tubulin polymerization assay

Porcine brain tubulin was prepared as previously described (Castoldi and Popov, 2003; Gell et al., 2011). GMPCPP, GTP, and ATP were obtained from Jena Bioscience. Anti-Biotin Antibody, PIPES, pluronic F127, D-glucose, DTT, casein, glucose oxidase, and catalase were purchased from Merck.

<u>Microtubule GMPCPP seeds</u> - 5 µl of biotin-labeled tubulin at 4 mg/ml (Cytoskeleton, Inc., unlabeled to biotin-labeled tubulin ratio - 150:1) was diluted in the 95 μl of the GMPCPP polymerization mixture (84 µl BRB80 (80 mM PIPES, 1 mM EGTA, 1 mM MgCl_2_, pH 6.8) with 10 μl of 10 mM GMPCPP and additional 1 μl of 100 mM MgCl_2_) and then pre-incubated for 5 min on ice as described previously (Gell et al., 2010). Microtubules were polymerized at 37°C for 1 h, centrifuged at 18000 x g with a tabletop centrifuge for 30 min at room temperature, and the pellet was subsequently resuspended in 150 µl BRB80. Microtubule seeds were stored at room temperature and used within one week.

<u>In vitro assay</u> - Flow chambers were assembled from 2 sizes of DDS-silanized coverslips (22x22 and 18x18 mm) and multiple parallel strips of parafilm between the coverslips to form separated chambers after a heat cycle on an electric heater. Flow chambers were then incubated for 5 min with 20 μg/ml anti-biotin antibody and subsequently blocked by 1% Pluronic F127 in PBS for 1 hour. Residual F127 was removed from the chambers by BRB80, and microtubules were attached to the surface by incubation of GMPCPP microtubule seeds in the chambers for 2 minutes. Unbound microtubules were removed by BRB80 that was subsequently exchanged by experimental buffer (BRB80 containing 10 mM DTT, 20 mM D-glucose, 0.1% Tween-20, 0.5 mg/ml casein, 1 mM Mg-ATP, 1 mM GTP, 0.22 mg/ml glucose oxidase and 20 µg/ml catalase). Microtubule growth was initiated by the addition of a mixture of proteins (10 µM FKBP12, 4 µM GFP-CRMP2A) in the presence of 12-16 μM tubulin (tubulin concentration was tested before each session to get similar tubulin dynamics).

<u>Imaging</u> - For fluorescence imaging, the total internal reflection (TIRF) mode of an inverted widefield microscope Nikon Eclipse Ti-E equipped with 100x HP Apo TIRF objective, H-TIRF module, LU-NV Laser Unit (all, Nikon, Tokyo, Japan), and sCMOS camera (ORCA 4.0 V2, Hamamatsu Photonics, Hamamatsu City, Japan) was used. Microtubule dynamics was visualized at the same microscope using interference reflection microscopy (Mahamdeh and Howard, 2019). Movies were acquired for 20 min with a 5 s time difference using NIS-Elements Advanced Research software v5.21 (Laboratory Imaging, Prague, Czechia). Analysis was done in FIJI using NanoJ-Core and Multi Kymograph plugins.

#### *In vitro* tubulin turbidity assay

Porcine brain tubulin (final concentration 25 µM), FLAG-tagged GFP-CRMP2A purified from HEK293T cells (final concentration 1.25 µM), and GST-FKBP12 purified from bacteria (final concentration 4.5 µM) were used for tubulin turbidity assay. To inhibit FKBP12, we used FK506 (final concentration 40 µM; InvivoGen, tlrl-fk5).

The assay was performed in 96well plates with glass bottom (Invitrogen, M33089) in a final volume of 100 µl. All reagents were mixed on ice in BRB80 buffer with 25% glycerol, 1 mM DTT, and 1 mM GTP), and the absorbance at 350 nm was measured every 15 seconds (for 1 hour, 37°C) using BioTek Cytation3.

#### EB3 plus-end tracking assay

IMCD3 cells were transduced with lentiviral particles carrying vector pTRIP overexpressing GFP-EB3 with or without overexpression of CRMP2A/CRMP2A S27A. For FKBP12 knockdown, a second lentiviral transduction with a pGhU6-mCherry vector carrying shRNA for FKBP12 was performed. Only cells holding both vectors (with mCherry and GFP signal) were selected. For FKBP12 overexpression, cells were transduced by pTRIP EB3 with or without overexpression of FKBP12. Transduced cells were then transfected by the mCherry-CRMP2A carrying vector. Only cells holding both vectors were captured. Live cell imaging was performed using a fluorescent microscope CARV II/Nikon Ti-E at 37°C, 5% CO_2_ (CFI Plan Apo VC 100X Oil, each 2s, for 1min). The data analysis was performed in FIJI software using the plugin MTrackJ. The mean velocity of the growing microtubule ends on the periphery of the cell with no contact with other cells was counted. The data are represented in the form of kymographs using the FIJI plugin MultipleKymograph.

#### Primary neurons cultivation, transfection, and immunocytochemistry

Primary cortical neurons from CRMP2A deficient (CRMP2A-KO) or WT embryos (E16.5-E17.5) were plated on coverslips (12 mm No. 1.5H; 0117520 – Paul Marienfeld) pretreated with Laminin and Poly-D-lysine (2 µg/ml and 50 µg/ml respectively) and cultured in Neurobasal media supplemented with B27 supplement, glutamine, and Penicillin/Streptomycin (ThermoFisher Scientific). Neuronal transfection was done at DIV1 using Lipofectamine 2000 in OptiMEM medium (ThermoFisher Scientific). Cells were fixed at DIV3-4 using 4% PFA and cold methanol unless stated otherwise. Cells were blocked with 0.1% BSA/PBS and incubated with primary (ON, 4°C), and secondary antibodies. Cells were visualized using a confocal scanning microscope Leica SP8 (PL FLUOTAR 25x/0.75 IMM or HC PL APO CS2 63x/1.40 OIL).

For correlation analysis, intensities of 100 pixels/cell (n=5) were measured and normalized (mean subtracted and divided by SD). Pearson correlation coefficient for CRMP2A and FKBP12/Pin1 signal was calculated. Differences between correlations were calculated according to Hypothesis tests for comparing correlations (Lenhard, 2014).

Microfluidic chambers were prepared from Sylgard 184 silicone (Dow) as described previously (Maimon et al., 2021) with modification of the wells prepared (4 x 7 mm diameter; Figure 5A). Neurons were plated on the “cell body part” of the chamber and four days after plating, neurons were stained using Cholera toxin Subunit B (ThermoFisher Scientific), and the “axonal part” of the chamber was captured for 40 hours (20-minute timeframe). Videos were then analyzed using FIJI plugin MTrackJ and average velocity was quantified.

#### Data analysis and presentation

GraphPad Prism was used for statistical analysis and graph illustration. Welch’s t-test or Mann-Whitney test was used to compare two independent groups according to the data normality unless stated otherwise. Adobe Illustrator and FIJI were used for figure preparation.

## Acknowledgments

This work was supported by The Czech Science Foundation grant (GACR 21-24571S), Charles University Grant Agency (GAUK 524218 for RW and GAUK 382621 for PB) and the project National Institute for Neurology Research (Programme EXCELES, ID Project No. LX22NPO5107) — Funded by the European Union–Next Generation EU.

We acknowledge the Imaging Methods Core Facility at BIOCEV, supported by the MEYS CR (LM2023050 Czech-BioImaging) and IPHYS Bioimaging Facility, supported by MEYS CR (Large RI Project LM2018129 Czech-BioImaging) and ERDF (project No. CZ.02.1.01/0.0/0.0/18_046/0016045).

## Author contributions

Conceptualization MB, RW, ZL; experiments performed by RW, DEC, JS, SB; method developed MB, JZ, RW, DEC, JS, PB; funding acquisition MB, RW, PB; resources and material MB, ZL, CJ, JZ, SB; supervision MB, ZL, CJ; figure graphics PB; writing RW, MB review & editing all authors.

