## Supplemental Figures for "Prolyl isomerase FKBP12 reduces axon growth and negatively regulates microtubule polymerization by inhibiting CRMP2A"

### Supplemental Files

#### Supplemental Figures

Figure S1

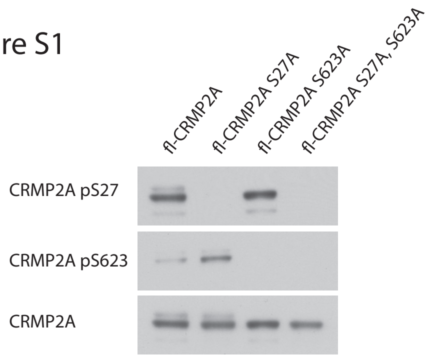


Figure S2

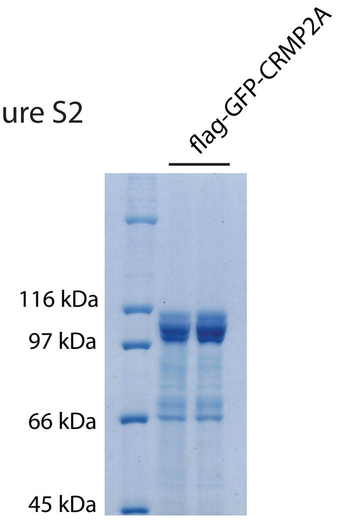


Figure S3

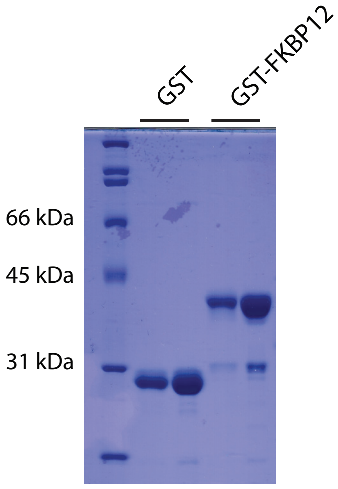


Figure S4

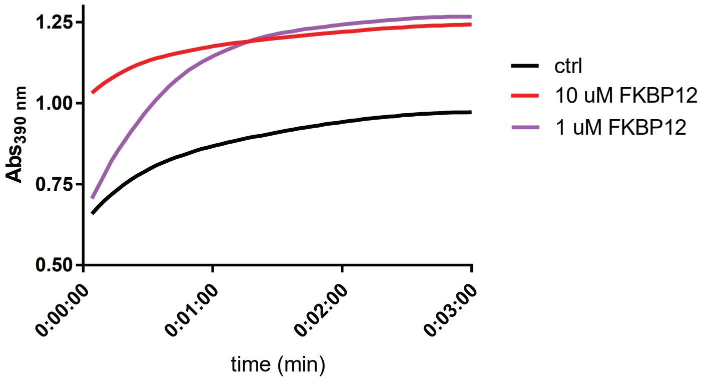


Figure S5 A

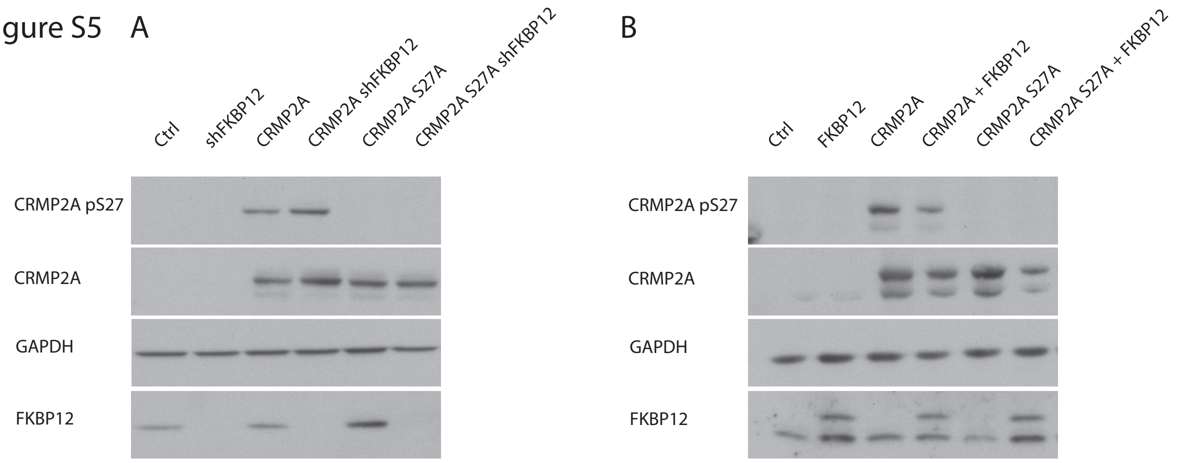

#### Supplemental Figure Legends

Figure S1. Dephospho-mimetic mutations used for pull-down experiments (Figure 2D-E). Dephospho-mimetic mutants are not recognized by anti-phospho CRMP2A specific antibodies, while they are recognized by anti-CRMP2A antibodies. Phospho-specific antibodies anti-CRMP2A-pS27 and anti-CRMP2-pS522 (corresponding to pS623 on CRMP2A) were used.

Figure S2. SDS-PAGE gel showing purified FLAG-GFP-CRMP2A protein used for microtubule polymerization assays stained with coomassie brilliant blue.

Figure S3. SDS-PAGE gel showing purified GST-FKBP12 and GST (used as a control) used for pull-down and microtubule polymerization assays stained with coomassie brilliant blue.

Figure S4. PPI-ase activity assay. Two different concentrations of FKBP12 (1 µM and 10 µM) were used to promote the digestion of a colorimetric substrate by chymotrypsin and to demonstrate PPI-ase activity of the purified FKBP12.

Figure S5. Western blots of cells used for GFP-EB3 assay, showing knockdown (A) or overexpression (B) of FKBP12 and overexpression of CRMP2A or CRMP2A S27A.
